## Supplementary Figure for "Cell culture-based shark karyotyping as a resource for chromosome-scale genome analysis"

### Supplementary Figures

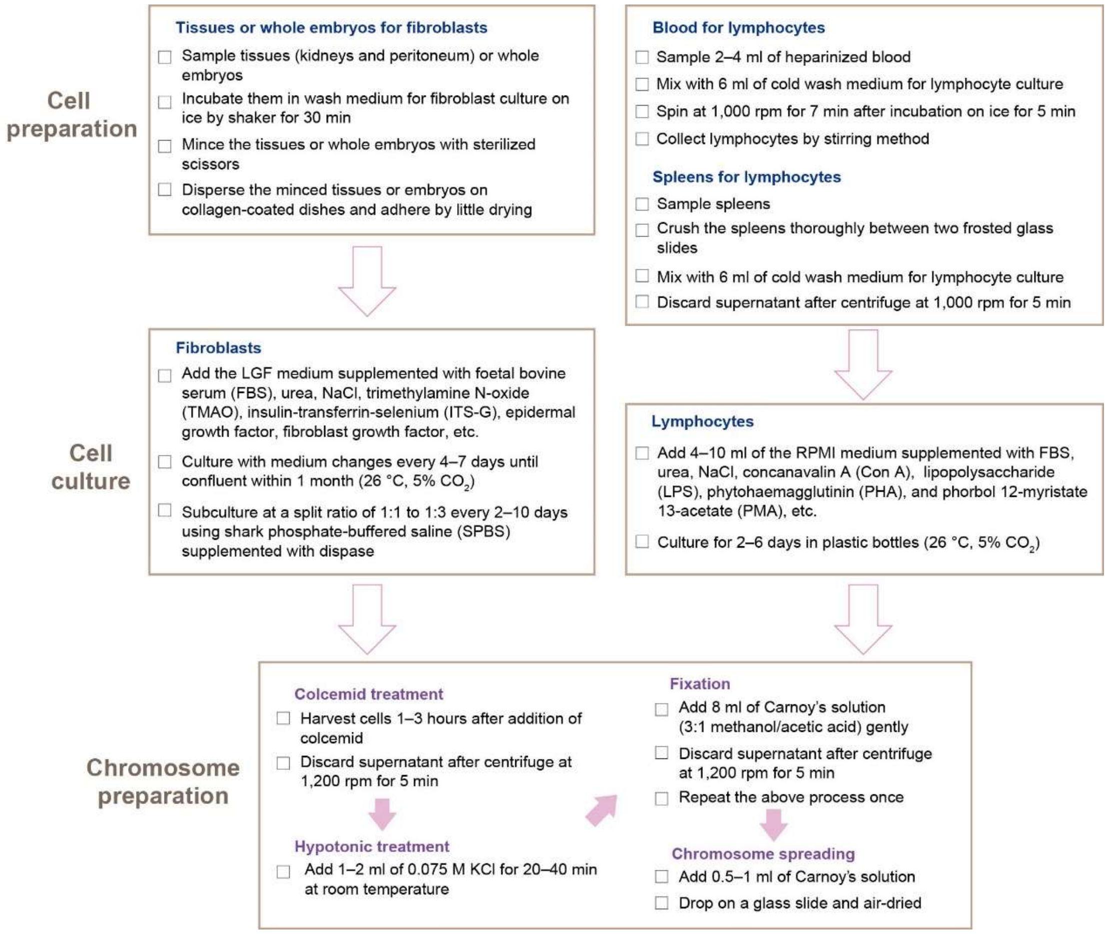


**Supplementary Fig. 1. Our cell culture protocol for shark chromosome preparation.** See Methods, Results and Supplementary Table 2 for the detail of medium formulations.

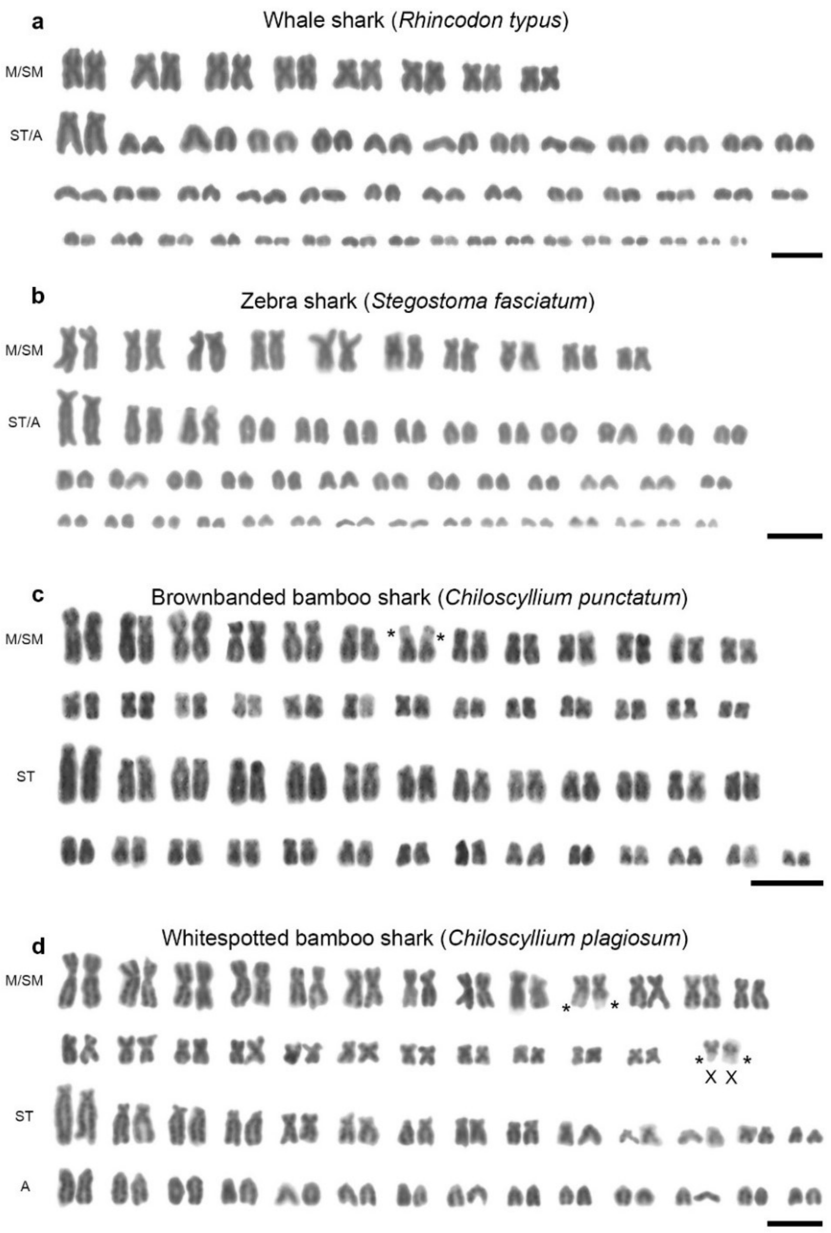


**Supplementary Fig. 2. Giemsa-stained karyotypes of female individuals. a** Karyotype of a female of the whale shark *Rhincodon typus* (2n = 102). **b** Karyotype of a female of the zebra shark *Stegostoma fasciatum* (2n = 102). **c** Karyotype of a female of the brownbanded bamboo shark *Chiloscyllium punctatum* (2n = 106). **d** Karyotype of a female of the whitespotted bamboo shark *C. plagiosum* (2n = 106). Asterisks indicate the positions of secondary constrictions. M, metacentric chromosomes; SM, submetacentric chromosomes; ST, subtelocentric chromosomes; A, acrocentric chromosomes. Scale bars show 10 µm. See Fig. 3 for male karyotypes and Supplementary Fig. 3 for metaphase spreads.

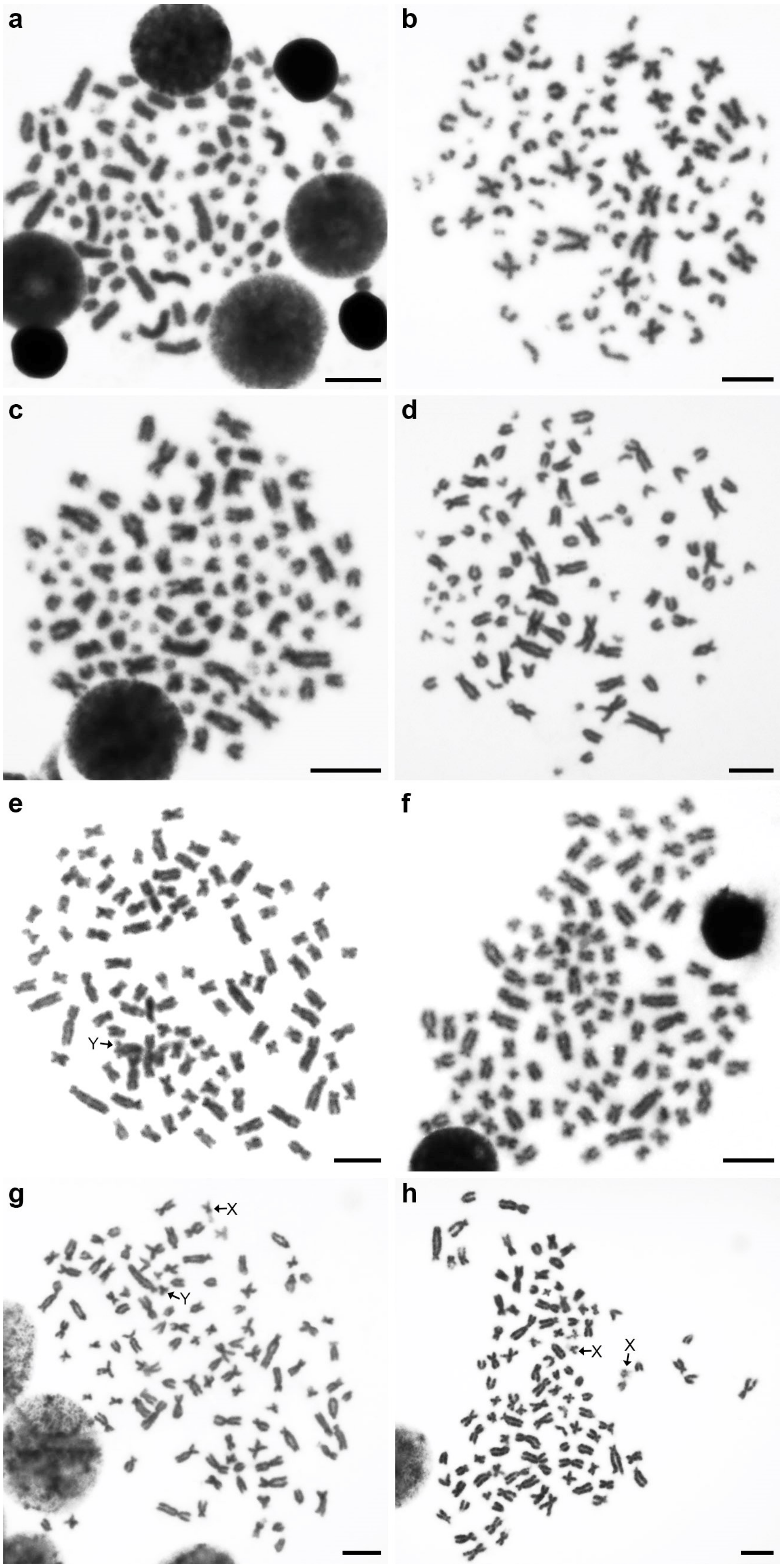
a
b
c
d
e
Y
f
g
X
Y
h
X
X

**Supplementary Fig. 3. Giemsa-stained chromosome metaphase spreads.** Metaphase spreads of males (**a, c, e, g**) and females (**b, d, f, h**) of the whale shark *Rhincodon typus* (2n = 102) (**a, b**), zebra shark *Stegostoma fasciatum* (2n = 102) (**c, d**), brownbanded bamboo shark *Chiloscyllium punctatum* (2n = 106) (**e, f**), and whitespotted bamboo shark *C. plagiosum* (2n = 106) (**g, h**). The ordered karyotypes from these metaphase spreads are presented in Fig. 3 (for males) and Supplementary Fig. 2 (for females). Arrows indicate putative sex chromosomes (**e, g, h**). Scale bars show 10 μm.

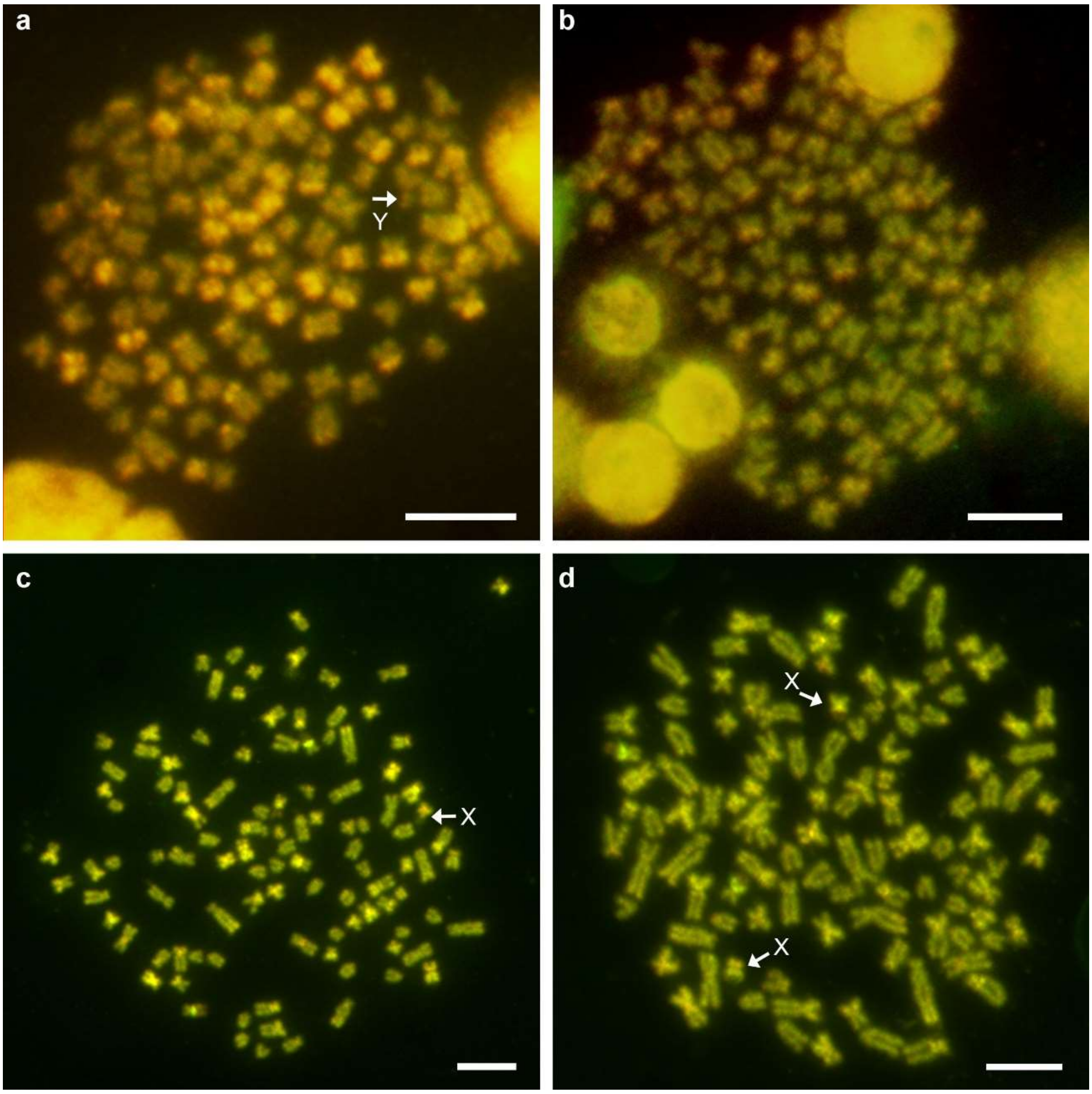


**Supplementary Fig. 4. Comparative genomic hybridization.** The FITC-labelled female genomic DNA (green) and Cy3-labelled male genomic DNA (red) on metaphase chromosome spreads are shown for a male (**a**) and a female (**b**) of the brownbanded bamboo shark as well as a male (**c**) and a female (**d**) of the whitespotted bamboo shark. Arrows indicate putative sex chromosomes. Scale bars show 10 µm.

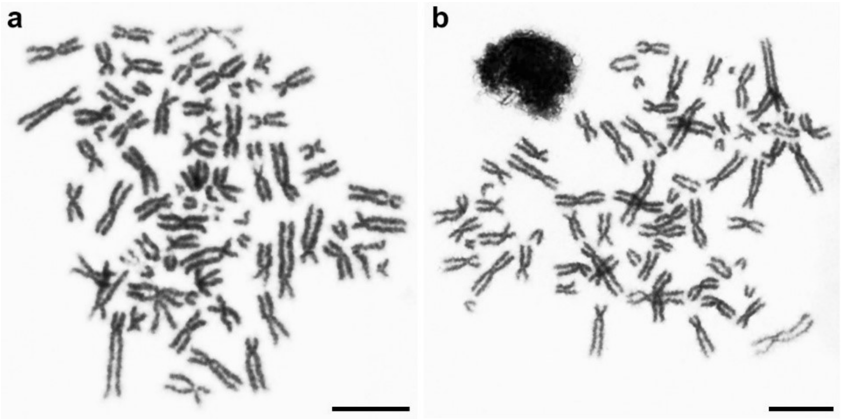


**Supplementary Fig. 5. Giemsa-stained chromosome metaphase spreads of two shark species in the order Carcharhiniformes.** We performed cell culture from tissues of two shark species in Carcharhiniformes, the banded houndshark *Triakis scyllium* and cloudy catshark *Scyliorhinus torazame* and karyotyping using the cultured cells. A 45 cm-long juvenile of the banded houndshark and an adult of the cloudy catshark were purchased from a commercial marine organism supplier in Mie Prefecture, Japan. After anesthetization with ice cooling, we used kidneys isolated from the banded houndshark for fibroblast culture and a spleen from the cloudy catshark for lymphocyte culture. The cultured cells of the banded houndshark and cloudy catshark were incubated at 26 °C and 20 °C in a humidified atmosphere of 5% $CO_2$, respectively. The other experimental conditions including medium formulations for cell culture and the method of chromosome preparation are described in the Methods of the main text. Metaphase spreads were obtained from the cultured cells from the banded houndshark (**a**) and cloudy catshark (**b**).These results confirmed their previously reported karyotypes[1,2]. Scale bars show 10 µm.
